# Sex-specific central regulation of reproductive activity revealed by the use of deslorelin in the grey mouse lemur, *Microcebus murinus*

**DOI:** 10.1101/2023.09.08.556849

**Authors:** Aude Noiret, Fabienne Aujard, Jeremy Terrien

## Abstract

Deslorelin is a GnRH agonist used in veterinary medicine to temporarily inhibit reproduction in domestic animals and is sometimes tested in captive species in zoo to control population or tame aggressive behaviours in males. However, some studies have revealed the inefficacy of deslorelin specifically in males, contrary to females that follow a classic long-term inhibition of the reproductive hypothalamic-pituitary axis through sexual steroid negative feedback. We implanted 5 males and 6 females grey mouse lemurs (*Microcebus murinus*), long-day breeders that display a complete inhibition of the reproductive system during winter, at the end of the short-day period, a few weeks before the breeding season. Contrary to females, which exhibited a classic inhibitory response to deslorelin, males testosterone levels increased as well as their testis size, which suggests a sex-specific sensitivity to the negative feedback of sexual steroids before the mating period. We propose that this sex-imbalance is related to the different life-history of males as opposed to females concerning reproductive tasks and behaviour.

## 1. Introduction

Seasonal breeders are known for their sex-specific reproductive axis regulations (Ball et Ketterson 2008). In many mammal species, males show early photorefractoriness to short day (SD) exposure, while females maintain an endogenous cycle which synchronizes with long day (LD) transition (Prendergast 2005; Perret et Aujard 2001). These features likely support males and females’ specific agendas regarding reproduction: while male prepare spermatogenesis in anticipation of the mating season, and engage in territory competition and courtship (Key et Ross 1999; Lane et al. 2010), females rely on a reserve of gametes from birth and delay reproductive activity until mating, gestation, lactation and young care (Czenze, Jonasson, et Willis 2017). During this time-shift where only males produce gametes, males and females’ sensitivity to the sexual hormones inhibitory action on the reproductive axis is likely to be different. Here, we investigated sex-specific responsiveness to sexual hormone negative feedback before the mating period by implanting grey mouse lemurs with a GnRH agonist, deslorelin (Padula 2005). As an agonist of GnRH, deslorelin primarily stimulates the release of sexual hormones in both males and females, which later translates into long-term inhibition of the hypothalamus-pituitary axis through a negative feedback (Trigg et al. 2006), resulting in a chemical sterilization. However, in some species, while deslorelin is efficient on females (Silvestre et al. 2009; Fontaine 2015; Goericke-Pesch et Wehrend 2012), males respond poorly (Eymann et al. 2007; Lincoln 1987; Aspden et al. 1998) with no modification of sexual activity. While the results in these species did not meet the expectations for an application in veterinary medicine (as population regulation, inhibition of aggressive behaviour), the sex-specific responsiveness to the molecule represents a good opportunity to discuss sex-specific reproductive regulation in an evolutionary point of view. As seasonal breeders, female *Microcebus murinus* show endogenous oestrous cycles that are synchronized with the LD photoperiodic exposure, while males enter in an early photorefractoriness after 14 weeks of SD exposure, which translates into rising testosterone levels and testis recrudescence (Perret et Aujard 2001; Terrien 2018). In a pilot study, a male was used to check on deslorelin efficacy and showed very low urinary testosterone concentration (7.8 ng.mg Creat.^-1^) compared to two non-implanted males (27.6 and 60.1 ng.mg Creat.^-1^) 15 weeks after implantation, despite large testis size. Here, we expanded this study with more animals, including females. For this, we implanted both sexes with deslorelin before males and females’ sexual reactivation, i.e. before the second half of winter to test for a sex difference in response to the increase in sexual hormones (testosterone and oestrogen). Despite originally looking for an inhibition of sexual activity in both sexes, we show instead contrasting results between males and females, and question this outcome in relation to the sex-specific life history of the species.

## 2. Materials and methods

### Animals and ethical concerns

Thirty grey mouse lemurs (*Microcebus murinus*), 14 males and 16 females all aged from 2 to 4 years and raised in good health in the breeding colony of Brunoy (MNHN, France, license approval n° E91-114-1), were included in the experiment. Temperature and humidity were maintained constant (24–26°C and 55%, respectively). The lemurs were fed with a fresh mixture (egg, concentrate milk, cereals, spicy bread, cream cheese, and water), banana, and were provided with *ad libitum* water. All described experimental procedures were approved by the Animal Welfare board of the UMR 7179, the Cuvier Ethics Committee for the Care and Use of Experimental Animals of the Muséum national d’Histoire naturelle, authorized by the Ministère de l’Enseignement Supérieur, de la Recherche et de l’Innovation (n°14075-2018031509218574) and complied with the European ethic regulations for the use of animals in biomedical research.

### Manipulation of the reproductive axis

Eleven animals (DIM deslorelin-implanted males, N=5; or DIF, deslorelin-implanted females, N=6) were implanted with a deslorelin implant (Suprelorin 9.4 mg) 6 weeks after short-day transition (SD: 10 hours light/14 hours dark), during early winter, at the time when both sexes reached an inactive sexual phase (Perret et Aujard 2001). Implants were applied on the interscapular zone under general anaesthesia (Alfaxalone 20 mg/kg IM + lidocaine injection at the implantation site). The implants were well supported, no itching lesions were observed and the wounds closed by themselves 2 days after the procedure. An additional surgical procedure was performed under general anaesthesia 50 to 35 days before LD transition for males and 1 month after transition to summer for females, in order to remove as much as implant as possible. At removal time, the implant was dissociated in several pieces. Although, the removal was complete and successful in most cases, some small implant pieces were left in the animals in some cases. In parallel, non-implanted animals (NIM non-implanted males, N=9; or NIF, non-implanted females, N=10) were also monitored.

### Assessment of reproductive activity

Sexual status was monitored from implantation until transition to long-days, i.e. summer season (20 weeks after implantation). Females were regularly checked by visual examination to determine whether they entered either proestrus or oestrus manifestation, which is easily observable in this species (Figure 1). In males, testis size (TS) was monitored each month (rated from 0 to 2 based on their size and consistency: 0= testes are up in the abdominal cavity and scrotal sacs are loose; 0.5= testes descend and measure 1 cm put together; 1= 2 cm; 1.5 = 3cm; 2= 4cm) (Figure 1). Sexual hormone levels (17-beta-Estradiol in females, plasma and urinary testosterone in males) were assayed in urine by ELISA after 2 months of implantation in all animals (“17-beta-Estradiol” in pg.ml^−1^, IBL, ref RE52041; “Testosterone” in ng.ml^−1^, Abcam, ref ab108666). Creatinine concentration (mg.ml^−1^) was used to normalize all urine measurements as an indicator of renal filtration activity (MicrovueTM Creatinine Elisa kit, Quidel R Corporation, ref 8009). Results are thus expressed in ng.mg Creat.^−1^ or pg.mg Creat.^−1^ for testosterone and 17-beta-Estradiol, respectively.

**Figure 1.**
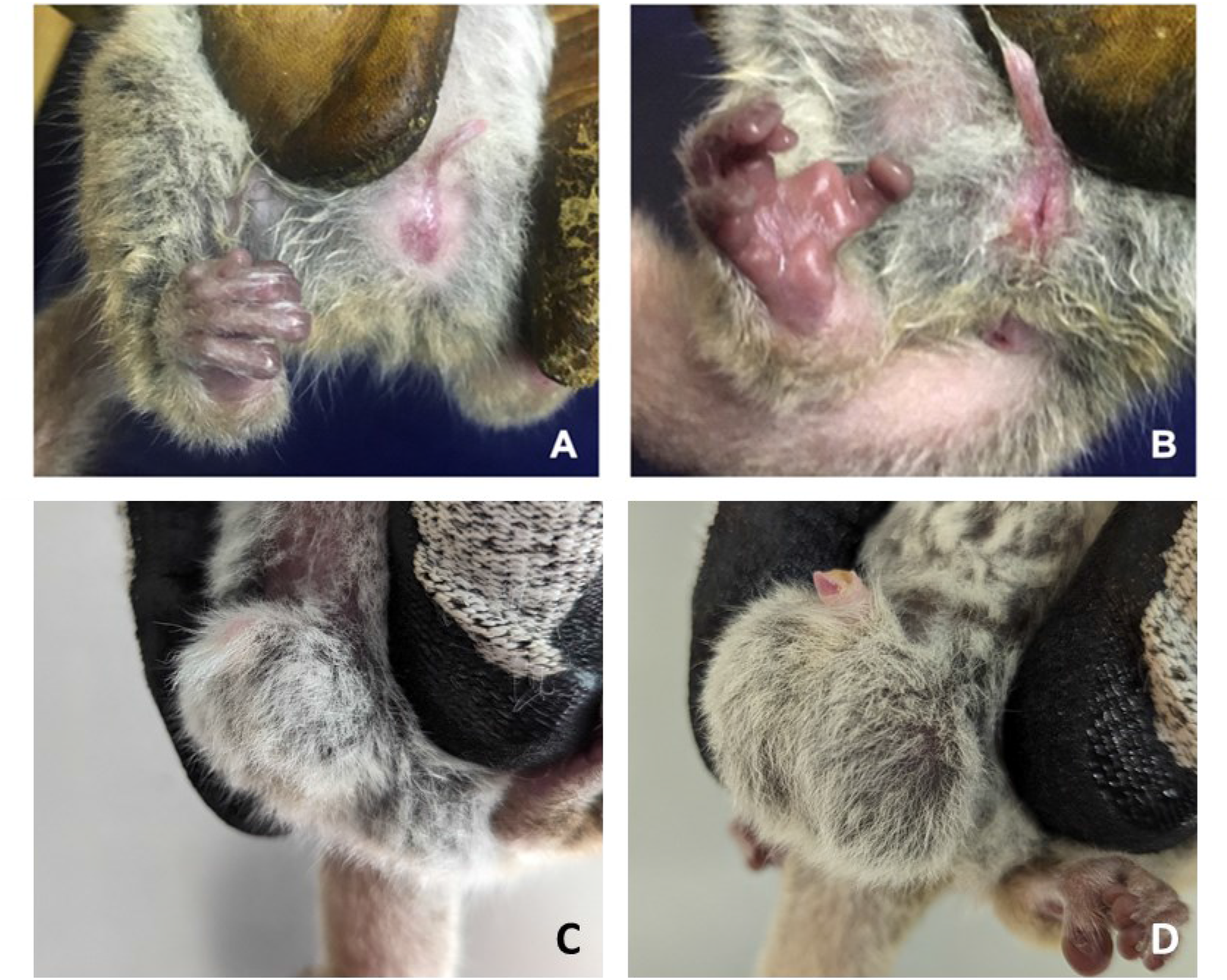
Photographs of the reproductive state in female (A and B) and male (C and D) mouse lemurs. Females implanted with deslorelin entering into (A) proestrus and (B) oestrus two weeks after implantation. Examples of males rated 0.5 (C) and 1.5 (D), according to testis size and consistency.

## 3. Results

### Effect of deslorelin implant in female mouse lemurs

All deslorelin-implanted females (DIF, N=6) showed early manifestation of reproductive axis activity between day 7 to 13 after implantation, i.e. 7 to 8 weeks after winter onset, as three females were in proestrus and 2 showed true open vaginas (Figure 1B). Females were checked 2 weeks after, and one still had a cicatricial oestrus. All females then entered into an inactive reproductive state (no oestrus) for the rest of winter, until photo-transition to LD. After LD transition, DIF did not show sexual reactivation, while estruses were observed in non-implanted females (NIF) about two weeks after LD transition (17 ± 3 days). We took off the implants 40 to 50 days after photo-transition to LD, and DIF showed variability in the timing of reactivation of reproductive axis (from 2 weeks to a month), which seemed to depend on the amount of deslorelin remaining under the skin after removal. Indeed, the implants broke-up into several pieces (up to 8), especially in females where they remained longer than in males, and some migrated far into the subcutaneous tissue. In four females, we only managed to extract ∼ 7 of the initial 9.4 mg (1.67 ± 0.23 mg either remaining or diffused) and they did not show reproductive activity at the same time than the other females which had the entire implant removed successfully (15 ± 2 days after the removal). One female with a remaining ∼1.5 mg implant entered oestrus 29 days after the removal.

The visual observations of sexual inhibition during winter in DIF were consistent with oestrogen concentrations in urine, which were significantly lower in DIF than in NIF (3034 ± 1112 ng.mgCreat^-1^ in DIF vs. 5910 ± 3328 ng.mgCreat^-1^ in NIF; W=11, p-value= 0.042 Figure 2A) 2 months after implantation (during the second half of winter).

**Figure 2.**
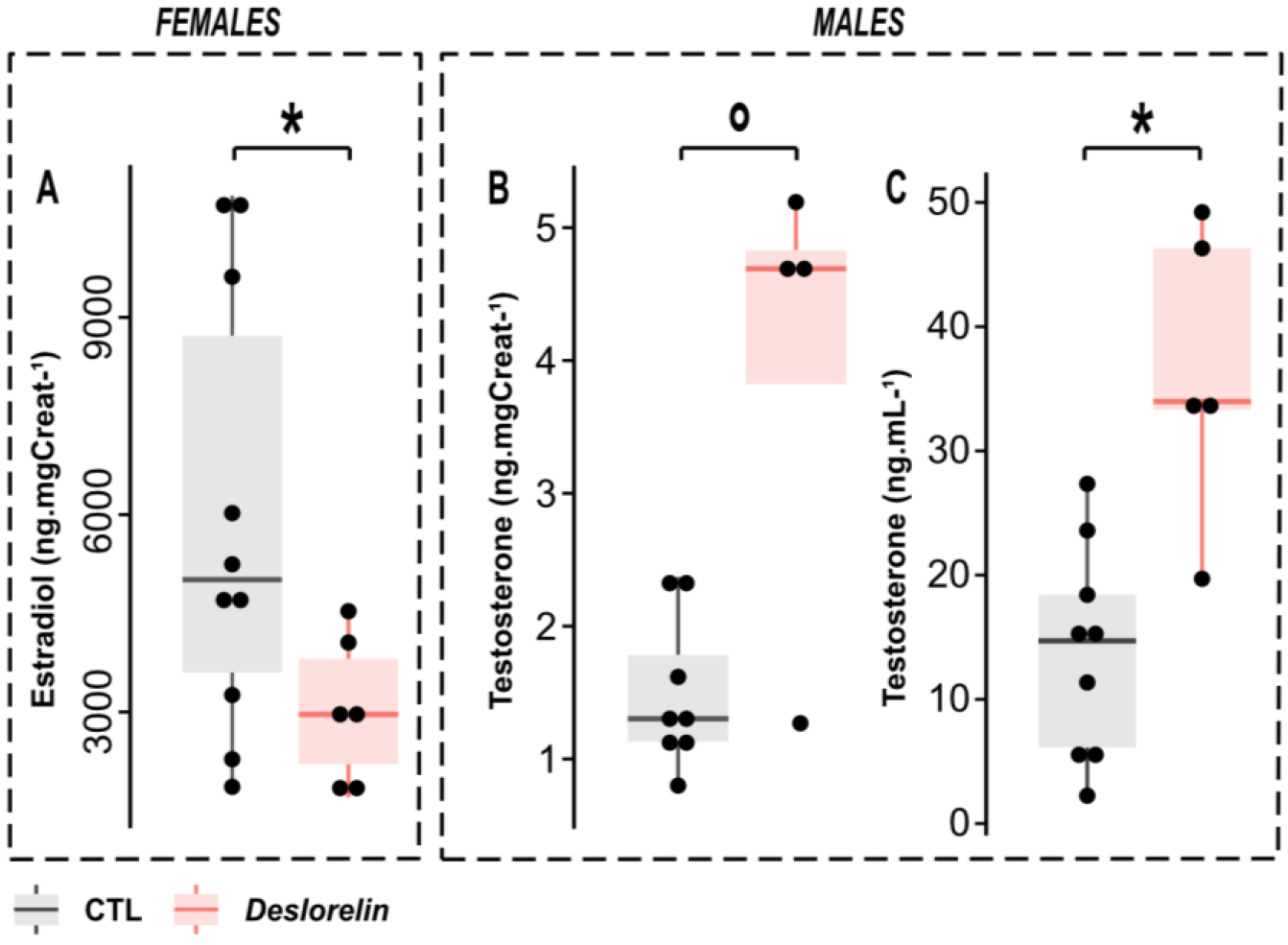
Sexual hormone levels in females (Estradiol) and males (Testosterone), in CTL animals (in grey) or after 2 months of deslorelin implantation (in red); A. Estradiol concentration in urine in females corrected by creatinine concentration (ng.mgCreat^-1^); B. Testosterone concentration in urine in males corrected by creatinine concentration (ng.mgCreat^-1^); C. Testosterone concentration in plasma in males (ng.mL^-1^). °: p-value < 0.1; *: p-value < 0.05.

### Effect of deslorelin implant in male mouse lemurs

In males, testis recrudescence began at week 9 after the beginning of the SD period, one month after deslorelin implantation (DIM: testis size was >0.5 by 10 ± 1.2 weeks after SD transition), while it started around week 14 for non-implanted males (NIM: testis size was >0.5 by 14 ± 2.2 weeks after SD transition). This earlier recrudescence in DIM was accompanied by a tendency for higher testosterone concentrations in urine until week 21 after SD transition (3.9 ± 1.8 in DIM vs 1.5 ± 0.6 in NIM; W=27, p-value= 0.073, Figure 2B). Plasma analysis significatively confirmed the augmentation of testosterone levels in deslorelin-implanted males compared to non-implanted animals (36.5 ± 11.8 ng.mL^-1^ in DIM vs 13.8 ± 8.5 ng.mL^-1^ in NIM; W= 43, p-value= 0.004, Figure 2C).

## 4. Discussion

As seasonal breeders, male grey mouse lemurs show earlier gonadal activity as compared to females, for they begin preparing sperm stocks long before the mating period. Moreover, this primate species presents specific reproductive life history traits (sperm competition, territoriality, exclusive maternal care), which adds to the gap in male-female biological and behavioural tasks regarding reproduction and young care. Hence, grey mouse lemurs are good candidates to study sex-specific central regulation of the reproductive axis and sensitivity to the sexual steroids negative feedback, using a GnRH agonist, deslorelin.

### The inhibitory effect of deslorelin is sex-specific in wintering mouse lemurs

In wintering, quiescent female mouse lemurs, the pattern of successive reactivation followed by inhibition of reproductive activity after deslorelin implantation is very similar to what is observed -and targeted-in domestic mammals such as cats, dogs, ferrets or dairy cows (Silvestre et al. 2009; Fontaine 2015; Goericke-Pesch et Wehrend 2012; Walter et al. 2011). Indeed, as a GnRH agonist, deslorelin triggers an initial upregulation of the reproductive axis, followed by a subsequent desensitization leading to a quiescent reproductive state. Given the effects observed in the pilot study (lowering of urinary testosterone despite large testes), we initially thought that the early recrudescence of testis size following deslorelin implantation was not synonym of testicular activity. Testis could be “frozen” in a big shape after an initial reactivation, showing no hormonal production, nor exocrine function, which ultimately meant that males developed desensitization of their gonadotropic cells, as it is described in dogs (Junaidi et al. 2003), and as we observed in female mouse lemurs. In our large-scaled manipulation however, we observed an opposite effect in male mouse lemurs, with an increase in testis size associated with the increase in urinary testosterone levels, suggesting the inefficiency of the negative feedback loop. Due to limited possibilities in blood sampling volume in those small animals, we did not assay for LH or FSH levels to acknowledge true desensitization of the gonadotropic axis, yet it is the general understanding about the action mechanism of GnRH agonists. In many species, males showed significant reduction of FSH and testosterone levels after deslorelin implantation, such as in dogs (Junaidi et al. 2003), eulemurs (Ferrie et al. 2011), as well as in baboons (Young 2013), cheetas, cats and ferrets (Bertschinger et al. 2006; Fontaine 2015; Goericke-Pesch et Wehrend 2012). Sex-specific effects of deslorelin exposure were previously described in the common brushtail possum, where females responded by a disruption of the normal estrous-cycle after an acute increase of LH, while males remained fertile after chronic deslorelin exposure and even sired as many offspring as the CTL males (Eymann et al. 2007). However in possums, testosterone and FSH levels decreased in response to deslorelin, which is yet another contradiction with our results. The male possums also lost the ability to respond to a surge of GnRH by an increased LH peak. Males of different species showed poor response to chronic GnRH agonist treatment, such as the marmoset monkey (Lunn et al. 1990; 1992), red deer stag (Lincoln 1987), tammar wallaby (Herbert et al. 2004) and bulls (D’Occhio et al. 2000). In bulls, LH release was associated with increased secretion of testosterone, persistent for the duration of deslorelin treatment (Aspden et al. 1998). It was suggested that bulls treated with GnRH agonist undergo the classical desensitisation of the pituitary and downregulation of endocrine function, but that other testicular factors could be involved in maintaining LH secretion, such as an increase in the rate of transcription and translation of LH β-subunit mRNA to LH (Aspden et al. 1998). Increased testosterone secretion in deslorelin-treated bulls was also associated with increased levels of testicular steroidogenic acute regulatory (StAR) protein and steroidogenic enzymes (Aspden et al. 1998). Whether this pattern is also implicated in the regulation of spermatogenesis in male mouse lemurs remains to be explored.

### Sex-specific response to deslorelin as a result of seasonal breeding and reproductive ecology

The time frame of deslorelin implantation is rarely discussed, although this parameter seems to be critical for deslorelin efficacy. In our study, implantation of deslorelin after mating could have maybe succeeded in downregulating sexual activity in males, when a decrease in energy allocation to sexual behaviour and spermatogenesis would not impact reproductive success. From the literature, it appears that deslorelin inefficacy is only observed in males, although the test is often made in the one sex - except for possums and marmosets. However in seasonal breeders, many studies have led to the observation that males and females don’t show the same regulation of the reproductive axis (Ball et Ketterson 2008). In winter (or SD), animals are sexually inactive (Simonneaux 2018), except for SD breeders as sheeps (O’Callaghan et al. 1992; Weems, Goodman, et Lehman 2015). However, while females start their reproductive cycle at the transition to LD, photorefractoriness to SD in males has been seen in many species leading to an earlier reactivation of the reproductive axis (Prendergast 2005; Ball et Ketterson 2008; Perret et Aujard 2001). This phenomenon allows to prepare sperm stocks, stimulate territorial and competitive behaviours, which in the end promotes reproductive success in a context of male-to-male competition (Key et Ross 1999; Lane et al. 2010). While we have relatively precise knowledge of the physiology of the reproductive hypothalamo-pituitary axis and its control, evolutive perspectives discussing its modulations across the different species and their life-history, which differs between sexes, are rarely evoked. Indeed, the feedback control of sexual steroids seems to emerge for different reasons in both sexes; in females, it regulates the estrous cycle, controlling the LH pulse and timing of ovulation in an ubiquitous fashion amongst mammals (Herbison 2020); in males it seems less stable amongst species and along life stages (Muller 2017), mitigating energy allocation to reproduction only when the conditions are favourable. Here, deslorelin implants were put after 6 weeks of SD exposure, when all the animals, males or females, were sexually inactive – when the gonads do not have any endocrine nor exocrine activity (Perret et Aujard 2001; Aslam et al. 2002). Females showed the expected acute responsiveness to GnRH agonist and subsequent negative feedback from sexual steroids. In males however, the increase in testosterone levels was not sufficient to inhibit the reproductive axis, and rather steadily boosted testis recrudescence until the mating window. Hence, not only the time-frame of energy allocation to reproduction is different between males and females, but also the efficacy of the negative feedback of sexual steroids. In the male horse, testosterone suppresses LH during the non-breeding season, while it enhances it early in the breeding-season (Irvine, Alexander, et Turner 1986). For males, higher testosterone levels translate into better reproductive success, which makes the variation of testosterone plasma levels a selective force for many reproductive behaviours specific to males, as stated by the challenge hypothesis (Muller 2017). Indeed, testosterone levels positively correlate with aggressiveness, territory defence, female fecundity, or even social instability of the alpha-male in a social group of primates (Muller 2017). Inversely, in some primate species that exhibit paternal care and aggressive behaviour simultaneously, males can be insensitive to testosterone during the reproductive period (Muller 2017). For the grey mouse lemur specifically, males engage in sperm competition, and display all the anatomic and behavioural features associated with this trait: high testis size/ body size ratio, penile spines, aggressiveness and territorial defence behaviours (Aslam et al. 2002; Wedell, Gage, et Parker 2002). In conclusion, it is not that surprizing that we obtained the exact opposite from the initial expected results in males - i.e. a boost of testosterone and testis size instead of a chronic inhibition of reproduction as seen in females. But whether the sex-specific response to deslorelin is due to testosterone desensitization of GnRH neuron cells in males or to other additional up-regulatory pathways remains to be demonstrated. However, this study allowed to further describe sex-specific reproductive regulation in a seasonal breeder and discuss another interesting result of sex-specific physiology in an evolutive point of view that would benefit from other similar studies in different species with various reproductive behaviours and ecology.

## Conclusion

Male and female grey mouse lemurs responded in opposite ways to deslorelin implantation after 6 weeks of SD exposure. Females showed signs of long-term inhibition of their reproductive axis, as commonly described in many species. Males on the other hand, showed advanced testis growth and increase in testosterone production, contrary to what was expected. This study proposes to use deslorelin to evidence sex-specific regulatory pathways of sexual activity in *Microcebus murinus* as a highly seasonal breeder and a species exhibiting sperm competition.

